# Lost dollars threaten research in public academic health centers

**DOI:** 10.1101/071860

**Authors:** Henry R. Bourne, Eric Vermillion

**Affiliations:** Authors are retired employees of the University of California, San Francisco. HRB was chair of Cellular and Molecular Pharmacology, and EV was Vice Chancellor— Finance

## Abstract

Decreasing federal and state support threaten long-term sustainability of research in publicly supported academic health centers. In weathering these financial threats, research at the University of California, San Francisco (UCSF) has undergone three substantial changes: (i) institutional salary support goes preferentially to senior faculty, while the young increasingly depend on grants; (ii) private and government support for research grows apace in clinical departments, but slowly declines in basic science departments; and (iii) research is judged more on its quantity (numbers of investigators and federal and private dollars) than on its goals, achievements, or scientific quality. We propose measures to alleviate these problems. Other large public academic health centers probably confront similar issues, but—except for UCSF—such centers have not been subjected to detailed public analysis.

---

Two grave financial threats endanger biomedical research conducted in publicly supported academic health centers: first, for more than two decades state governments have steadily reduced their previously generous support for higher education; second, five decades of large annual increases in the National Institutes of Health (NIH) budget ended abruptly in 2004 and will not soon resume, owing to a stagnant economy and political gridlock. One leading center, the University of California San Francisco (UCSF), has weathered these twin threats financially, but— based on a recent detailed analysis (1)—not without substantial risks to the quality, direction, and sustainability of research.

It may appear surprising to mention risks in relation to UCSF, whose NIH grant portfolio, record of discoveries, and research awards rank it as one of the biggest and most successful public academic health centers in the US. The surprise probably reflects a lack of prior detailed analyses of research funding in large public academic health centers, which are likely to share many of the financial challenges and coping strategies observed (1) at UCSF. If so, the following account will interest a wide spectrum of researchers, planners, academic leaders, and research funders.

## Financing UCSF’s research: overview

In the 30 years from 1984 to 2014, UCSF’s annual revenues—including those of its clinical enterprise and the four schools of its academic Campus (Dentistry, Medicine, Nursing, and Pharmacy)—grew 10-fold in nominal dollars, from $450M to $4.45B (1); in inflation-corrected dollars (2), the increase was 3.4-fold. Relative contributions of different revenue sources during this period (Figure 1, left panel; all corrected for inflation; 2) reveal changes that profoundly altered character and direction of UCSF’s research. A six-fold clinical expansion of UCSF’s clinical revenues mirrored the 5.5-fold growth of the US medical care industry over the same period (1). UCSF’s clinical enterprise garnered only 30% of its total annual revenue in 1984, but 54% in 2014 (1). In contrast, state support in 1984 accounted for 22% of total revenue, but only 4.4% in 2014—a decline that reflected a 31% decrease in UCSF’s annual state appropriation between 1984 and 2014, paralleling a nation-wide decline of state support for higher education. Research contracts and grants increased 3.5-fold, but provided similar proportions of revenue (27 *vs*. 26%) in both 1984 and 2014.

**Figure 1.**
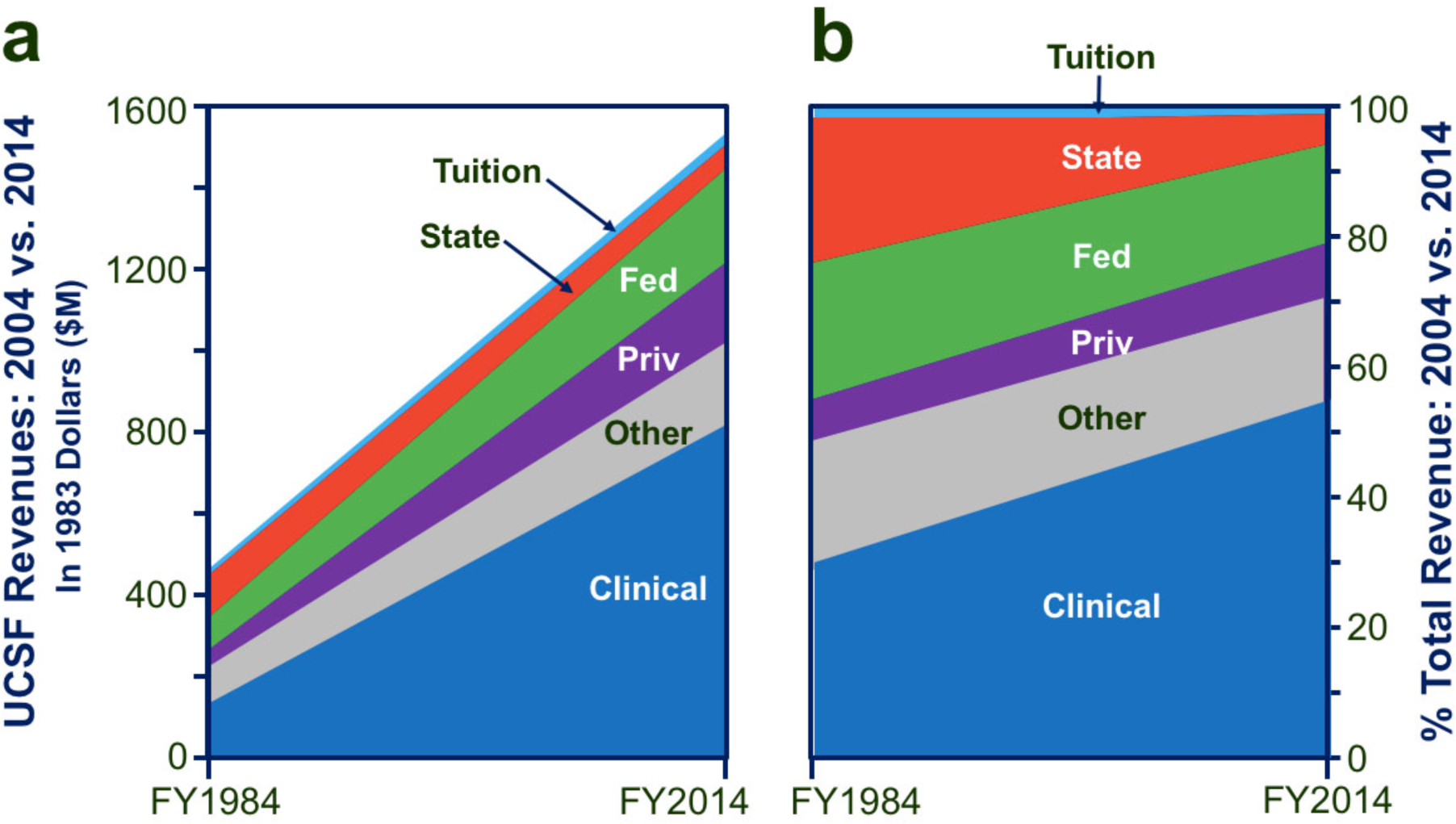
Comparison of UCSF revenues in fiscal years 1984 *vs*. 2014. Panels a and b show changes in this period for (top to bottom, in each panel): tuition (light blue), state support [State, officially the “State Educational Appropriation” (1); red], federal contracts and grants (Fed, green), private contracts and grants (Priv, purple), other (including miscellaneous revenues unrelated to research, gray), and the clinical enterprise (darker blue). a. revenues in millions of dollars (1983 dollars, corrected for inflation). b. revenues as a percentage of total UCSF revenues. Panels are simplified versions of Figures 1-4b and 1-4c, p 15, in ref. 1 (pp 14 and 15).

These changes exerted significant effects (1): (i) loss of state support progressively eroded UCSF’s ability to pay faculty researchers’ salaries, initiate new programs, recruit first-class scientists, and reward creative scientists; (ii) growth of the clinical mission converted an academic enterprise focused on research into an expanding jumble of hospitals and health care deliverers attached to a relatively shrinking academic appendage; (iii) remarkably, however, by reshaping its research goals, guidance, training, financing, and administration, UCSF attracted increasing research support from external funders.

Less easily quantitated changes have made it even harder to sustain high-quality research (1). Many faculty researchers perceive UCSF’s leaders—compared to their predecessors 20 years earlier—as focusing most of their attention on competing for patients and clinicians and judging clinical and research success primarily in dollars and national ranking, rather than in terms of goals or quality of efforts to achieve them. Nationally, citizens and legislatures now urge cuts in academic research dollars, asserting that research can pay its own way, presumably in patent income or support by industry. Such views ignore the fact that research efforts, fundamental or applied, necessarily fail before they finally succeed. This fact is the core rationale for government to sustain research as an essential public good.

## Decreased state dollars: soft-money salaries

Loss of almost a third of its state support (1) has gravely damaged UCSF’s ability to contribute to researchers’ salaries. Until the 1990s, state dollars bountifully supported salaries of faculty and administrative staff involved in research and its funding. Tenure-track faculty received a “Full-Time-Equivalent” salary, which covered ∼75% of total salary for most faculty in basic science departments and 30-60% for tenure-track faculty in clinical departments. In 1984, UCSF researchers could feel sure their success or failure would depend on quality of their research, rather than on available research funds or salary. Now such confidence is rare, because the reduction in the state educational appropriation deprives UCSF of at least $90M (2014 dollars; 2) every year (1,3). This appropriation was designed to pay “full-time-equivalent” salaries for tenure-track faculty, plus administrative support for research and teaching. Today that lost $90M could pay all grant-derived salary of all tenure-track faculty researchers almost four times over, leaving a large surplus for research administration and other academic functions (4). But the $90M (2% of total UCSF revenues; 1) cannot be replaced by student tuition (5), grants, or clinical income— leaving a single viable option: philanthropy.

UCSF’s dependence on grant dollars for faculty researcher salaries generates difficulties well beyond the the reduced state educational appropriation. In a city beset by soaring increases in housing and living costs (1), 62% of the 1,498 faculty who receive any salary from grants receive no state-derived instructional salary dollars whatever (Figure 2). Four of every five faculty within this 62% are non-tenure-track members of clinical departments (1); their grants pay part of their salary, and most of the rest comes from clinical care. The number (and proportion) of clinical department researchers highly dependent on grants for salary has grown immensely, because clinical expansion since 1984 outpaced Campus growth 3-fold (Figure 1), and non-tenure-track faculty in clinical departments receive few or no state salary dollars.

**Figure 2.**
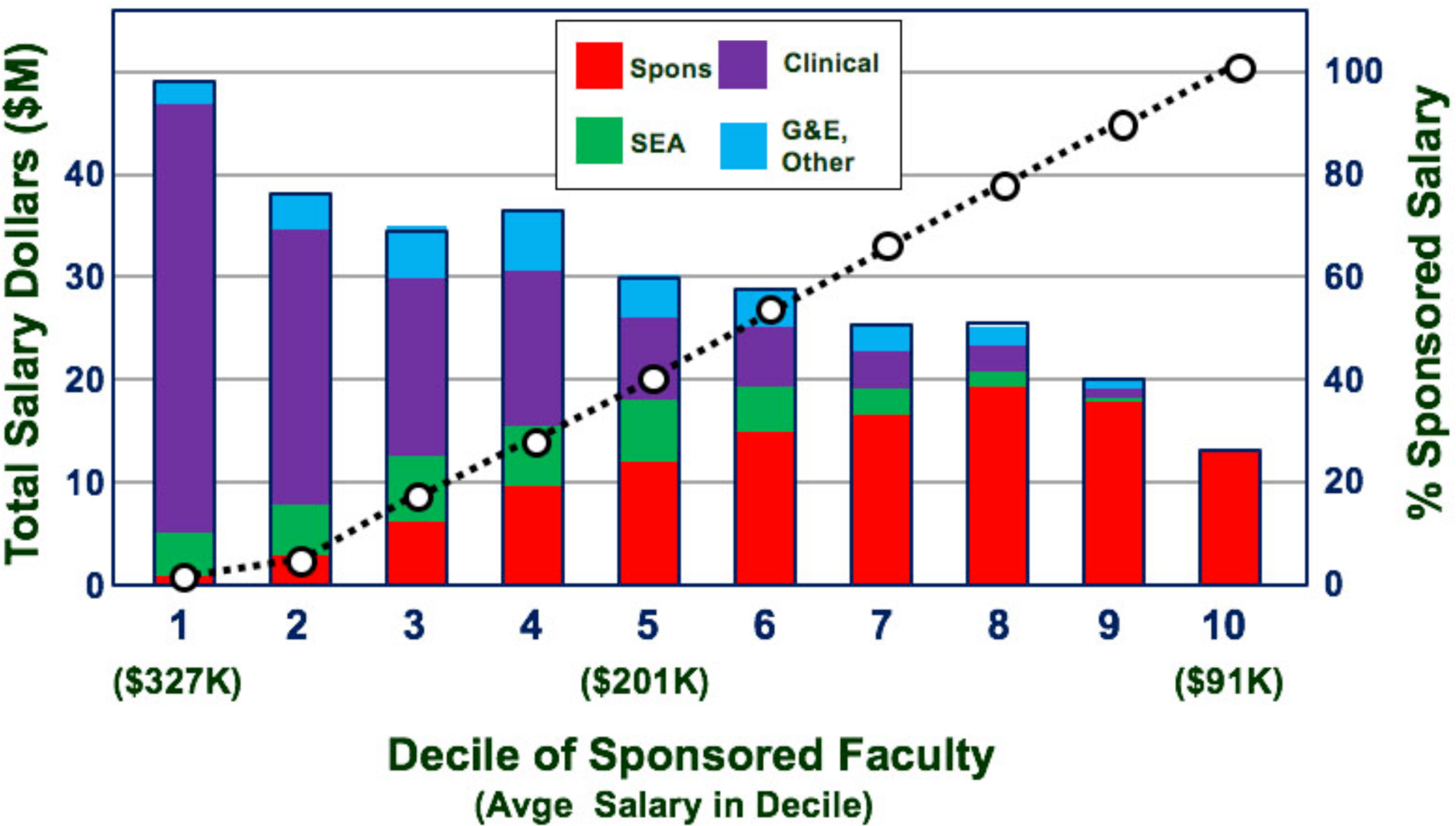
Salary sources in 2014 for 1,498 research faculty who received at least some portion of their salary from sponsored research. Columns represent 10 deciles of faculty (either 149 or 150 individuals, numbered 1-10 from left to right), ranked in ascending order of proportion of salary coming from sponsored sources. Each column shows millions of dollars (numbered on the left-hand ordinate) for an entire decile, by source (bottom to top): sponsored (red); state support (green); professional fees (clinical, purple); gifts and endowments and other (G&E, other). White circles, connected by a dotted line, represent the % of sponsored salary in each decile (right-hand ordinate). Dollar numbers in parentheses below decile numbers at the bottom of the chart show total salary (thousands of dollars, K) for average individual faculty in deciles 1, 5, and 10. The chart is a simplified version of Figure 6-1 in ref. 1 (p 93).

Figure 2 shows sources of salary dollars for UCSF’s 1,498 faculty with grant-supported salary (59% of all UCSF faculty) who derive at least some income from “sponsored” sources (i.e., grants). Ranking all 1,498 “sponsored” faculty in deciles by proportion of sponsored salary reveals disturbing surprises. Total annual salaries for those with the lowest proportions of sponsored salary (4% or less; deciles 1 and 2), are three-fold higher than salaries of those who earn 90-100% from sponsored dollars (deciles 9 and 10). Ironically for a university proud of its research mission, its lowest-paid (and younger; 1) sponsored faculty devote all their effort to research and receive no state salary dollars (green segments in the figure), while the best-paid in this group—older, focused on clinical care more than on research in deciles 1-3 received 45% of all state dollars paid to sponsored faculty, but only 8.6% of sponsored salary (green *vs*. red segments, respectively).

Most sponsored faculty depend on grant dollars more than is commonly thought. Leaders like to say that sponsored salary, averaged over all 1,498 sponsored faculty, averages less than 38% of total salary. But if we eliminate the best-paid 30% of these faculty (with average grant salary less than 10% of total salary), we calculate that the remaining 1,050 sponsored faculty depend on grants for (on average) 65% of their total salary, and thus are highly vulnerable to loss of any fraction of grant support. The high quantitative prevalence of soft-money salary is masked by giving more state-derived salary dollars to well-paid senior faculty, who are minimally involved in research, than to junior researchers.

We should worry about soft-money salaries for four reasons. (i) Paying ever-higher fractions of salary from grants often requires researchers, especially in early years of their careers, to obtain grant support for multiple separate projects and thus devote precious time to grant applications rather than focus intensely on a single important project. (ii) Formerly, hard-money salaries elevated the institution’s stake in its investigators’ success and fostered their loyalty to the institution and one another, making research more than a way to pay rent for laboratory space; today’s soft-money salaries do neither. (iii) Growing numbers of grant applications make soft money increasingly hard to get (6). (iv) Although the search for new knowledge always risks failure, scientists who know they can afford their families’ next meal can tackle harder, more rewarding scientific questions, like those addressed at UCSF in the 1970s through the 1990s (1).

## Sustainability and investment: bricks and mortar *vs*. people

Dwindling state dollars and stagnant federal dollars strain UCSF’s ability to sustain its research enterprise. To compensate for lost external support, publicly supported academic health centers increase the portion of faculty salary paid from grants (see above), maximize federal reimbursement of indirect costs of research, issue debt to build new research facilities, and solicit gifts and endowments. While these strategies help pay UCSF’s research bills, they also combine to create risks for research.

*Maximizing reimbursement of indirect costs*. In fiscal 2014, external private and federal research funders paid UCSF $992M, 80% of which were direct costs of research under investigators’ supervision. The remaining 20% represented partial reimbursement to the institution by funding agencies of the indirect costs of facilities and administration required to support the research—dollars commonly termed “indirect cost recovery” (ICR). In 2014 UCSF’s ICR amounted to $194M, or nearly 10% of that year’s non-clinical (Campus) revenue. Most ICR dollars came from NIH, calculated by complex formulas from the estimated facilities and administration costs UCSF incurred to support research (7). Distributed by the Chancellor, ICR dollars represent UCSF’s strongest and most flexible tool for shaping the research enterprise. Moreover, reliable flow of ICR—which assures bond holders that the university will repay them—has enabled the University of California (and thus UCSF) to sell large construction bonds since the 1980s.

ICR dollars also shape modes of payment and actual conduct of research in unintended ways. One is obvious: federal rules specify that indirect costs are reimbursed in direction proportion to direct costs incurred. This creates a perverse incentive for institutions to maximize payment of faculty salaries as direct costs of grants (8). Lacking hard state dollars to pay annual sponsored faculty salaries and benefits ($148M in 2014), UCSF requires its researchers to obtain that sum as direct costs on grants. By federal rules, this direct cost salary earns an ICR bonus of $60M. Endowments and state funds will never provide funds equivalent to the direct costs, the ICR, or their sum, which in 2014 was $208M (9).

*Construction and debt*. By 2015, UCSF had built a large new campus near Mission Bay in San Francisco, mostly devoted to research, but also including housing, recreation and parking space. Of the cumulative cost, $1.59B, philanthropic gifts have paid only 24%, while incurred debt accounts for 61% (47% paid from sale of UC bonds, plus 14% in a capital lease). UCSF pays this debt’s principal and interest by tapping ICR income for 38% of the costs, housing and parking income for 9%, equity (cash on hand) for 11%; and state funds for only 4.8% (1).

In 2015, UCSF owed $2.33B, distributed between its clinical enterprise (38%) and Campus (62%) (1); debt service (paying principal and interest) cost the hospital $43M, the Campus $59M. Here we encounter a second potentially perverse incentive (8): in 2014 UCSF received $45M in federal ICR (23% of its total ICR) as reimbursement for interest on loans for constructing research facilities and depreciation of those facilities; this covered much of the Campus’s annual debt service (10). New building and other needs will drive its debt and debt service even higher: by 2025, debt service is projected to more than double (to $160M; 1).

Compared to present annual Campus revenues—$2.06B in 2014, probably higher in 2025—that $160M seems modest, especially since ICR will reduce it further (1). But the sanguine view of debt service as a valuable source of ICR dollars is not quite right, because research expansion can create quantity at the expense of quality: debt-based expansion of research facilities absorbs university dollars that could otherwise be used to support research in existing facilities, reduce soft-money salary, explore exciting questions before external funds arrive, and improve research training. During a research building’s first decade, administrative costs, maintenance, utilities, and populating the building with new faculty far exceed federal ICR derived from both debt service and direct research costs charged in the new facility (1).

*Gifts and endowments (G&E)*. UCSF’s endowments totaled $1.99B in 2014, with a $68.3M net payout (1). These endowments help fragments of the Campus, because individual faculty, departments, and smaller administrative units control most endowment dollars, as specified by myriad donors. Of about 2,000 separate endowments, only 67 are big enough to yield annual payout comparable to a $200,000 faculty salary, and only five yield more than a $1M annual payout per endowment (1). Such endowments cannot be used to change overall quality and direction of UCSF’s research mission. Similarly, in 2014 the institution attracted $445M in some 31,000 non-endowment gifts, mostly to help pay for a building, study a disease, or for other circumscribed purposes that do not contribute generally to programs, productivity, stability, or sustainability of the Campus or Schools.

*Risks of UCSF’s funding strategies*. While all these financial strategies to fund research furnish genuine benefits, separately and in combination they also increase risks associated with the scarcity and arbitrary quality of government and private funding. Much of the increased risk comes from incentives to maximize ICR by increasing soft-money faculty salaries and issuing debt to build new research facilities, which together accounted for 51% of all ICR in 2014 (1). Such risks are rooted in one fact: all UCSF’s research funding strategies aim to maximize quantities of external funds available for research and minimize costs paid by the institution. It is not “perverse” for UCSF—deprived of former state funding, unable to offset state cuts by increasing tuition revenue (5), and facing fierce competition for researchers and research dollars—to take advantage of federal rules; not to do so would be folly. But those rules create perverse incentives to expand research quantity (of facilities and more researchers) in institutions eager to replace lost state dollars with grants and ICR. Diverting attention from research goals and quality allows quantity to pre-empt quality. As dollars from external sources become scarcer, pre-emptive risks grow.

Here we highlight additional risks:

1. *Out-sourcing evaluation of researchers.*Soft-money salaries (sources of 30% of annual ICR) tempt UCSF to defer to external funders and donors as ultimate judges of research quality. Today’s leaders lack the time and expertise to judge young scientists, says one prominent leader, who prefers letting NIH grant reviewers winnow researchers by “Darwinian selection” (1). This approach is risky, because it disproportionately subjects beginning scientists to NIH funding decisions that are arbitrary and inaccurate (owing to the high ratio of good grant applications to grant awards). In contrast, by enlisting chairs and faculty in evaluating researchers, earlier Campus leaders helped UCSF to select and retain top-flight academic scientists who consequently proved loyal to the institution— and *vice versa*.
2. *Providing institutional support to departments based on ratio of ICR dollars to square feet of research space.* UCSF recently announced this ratio as its sole quantitative criterion for furnishing institutional support to departments, in regulations that did not mention research quality (1). Quantitative guidance is necessary, but using ICR as a primary criterion risks supporting mediocrity.
3. *Favoring established over young scientists.* UCSF disproportionately pays institutional salary dollars to established older researchers, who could also easily earn more soft-money salary. Modest shifts toward supporting young scientists who need it will improve research quality; larger shifts will do even better.
4. *Preferential treatment of older researchers, job insecurity, and arbitrary grant reviews damp creativity and innovation*.
5. *Expansion of research facilities can divert dollars from high-quality science.* New buildings and new debt take resources from researchers already at the institution.
6. *Predominant targeting of gifts or endowments to faculty and departments* shifts responsibility for quality from an institution’s researchers to parochial and potentially arbitrary judgments by donors, administrators, and individuals.

## Changing research in different department types

How do changing patterns of external research funding affect researchers and the conduct of research? In UCSF’s School of Medicine, external research funding from 2004 to 2014 flowed to basic science departments, clinical departments, and Organized Research Units, or ORUs (Table 1). This ten-year period began just as the rapid annual growth of federal research dollars suddenly stopped (1,6), and followed an earlier decade of falling state support for UCSF. During the latter period (2004-2014), NIH budgets rose slightly in nominal dollars, but plummeted by 21% in inflation-corrected 2004 dollars (1,2). In the same inflation-corrected dollars, the School’s external research funds (government and private) rose by 16%, driven by a 31% boost in clinical department research dollars; research dollars fell 10% in basic science departments and 16% in ORUs (Figure 3). Research revenues grew faster in clinical departments than in non-clinical departments, because business and the public value translational research over fundamental research and clinical projects attract more private research dollars. Private dollars rose 57% in clinical departments and 29% in ORUs, but fell 18% in basic science departments. Federal funds rose 18% in clinical departments, but fell 21% in ORUs and 10% in basic science departments (1).

**Table 1.** Research in Department Types (2014): Basic Science, Clinical, and ORUs^*§^.

|  |  | **Basic** | **Clinical** | **ORU** | **Total** |
| --- | --- | --- | --- | --- | --- |
| **Faculty with grant support** |  | 112 | 1223 | 73 | 1,408 |
| **Research contracts & grants** |  |  |  |  |  |
| **Federal** | **\$M** | 82 | 389 | 83 | 553 |
| **Non-federal (private)** | **\$M** | 17 | 174 | 23 | 214 |
| **Federal + non-federal** | **\$M** | 99 | 563 | 106 | 769 |
| **Research C&G per faculty member** | **\$K** | 884 | 460 | 1,452 | 546 |

\*Confined to sponsored faculty and research dollars in UCSF's School of Medicine (SOM), this table omits the small amounts of research support derived from state or local governments. The SOM contains nearly all of UCSF's clinical faculty who receive some salary support from research grants, and its research contract and grants represent ~83% of UCSF research funding. Its nine basic science departments include seven traditional departments whose research is primarily focused on biological investigation in wet labs, plus Departments of Epidemiology and Biostatistics and Anthropology, History, and Social Medicine. The SOM's 20 clinical departments teach students and resident physicians and take care of patients; their faculty engage in wet-lab or dry-lab research, or sometime both. Six ORUs are charged with responsibility for specified subsets of research goals (e.g., understanding and treating cardiovascular disease or cancer); their members also have appointments in other Departments; their research focuses on both wet-lab biological studies dry-lab investigation of patient populations, disease, and different kinds of patient management and therapy.
§Data for numbers of faculty with grant support and dollars of research support are found in ref. 1, in (respectively) Table 8-2 (p 118) and Table 9-2 (p 139).

**Figure 3.**
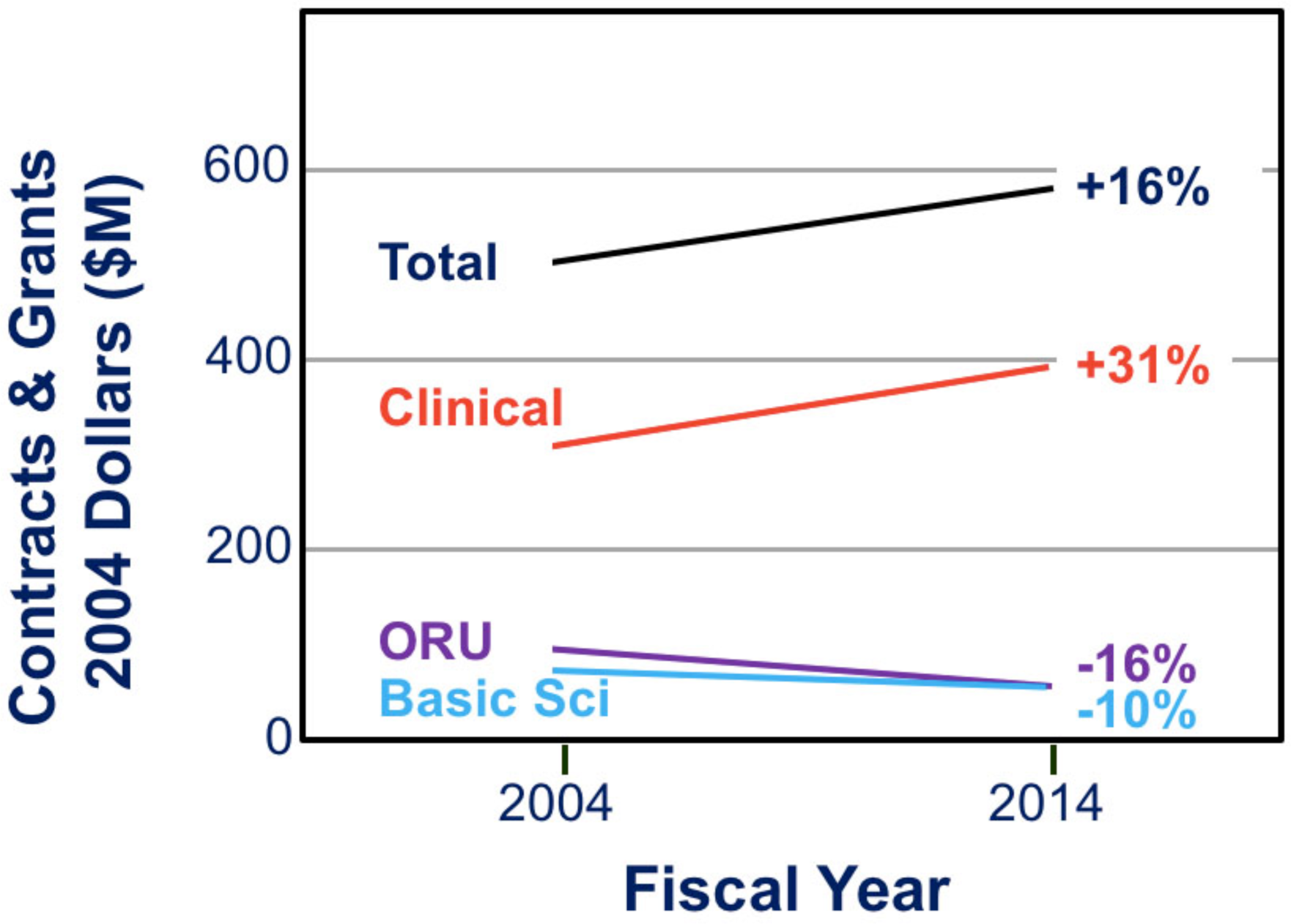
Changes in sponsored research funds received by different department types in UCSF’s School of Medicine (SOM) in 2014 *vs*. FY2004. Ordinate: millions of dollars (in inflation-corrected 2004 dollars). Abscissa: two fiscal years are shown Lines of different colors connect (from top to bottom): total sponsored SOM dollars, black; clinical department dollars (red); ORU dollars (purple); Basic science department dollars (blue). Corresponding percentage values (right) indicate the % change of that department type’s dollars from 2004 to 2014. The chart is a simplified version of part of Figure 7-1 in ref. 1 (p 108).

The most influential driver of clinical *vs*. non-clinical research funding is simple: expansion of UCSF’s clinical enterprise (Figure 1) allowed clinical departments to hire more sponsored faculty MDs and support part of their salaries from clinical income (11); in contrast, non-clinical faculty did not increase because ever-shrinking state support made it harder to support PhDs, whose salaries consequently became more dependent on grants. While basic science departments focused on wet-lab biological research, clinical departments increasingly shifted to dry-lab patient studies, probably because dry-lab approaches require shorter research training (attractive to MDs), less equipment and start-up costs, and less specialized lab space.

These changes (Figure 3) help to explain the differing feelings of clinical *vs*. non-clinical researchers about their research prospects (1). While established senior-level clinician-scientists feel sanguine about the future, senior basic science department faculty (unless they enjoy generous support from the Howard Hughes Medical Institute) often express discouragement about the future of fundamental research and even more about their departments’ weakened ability to compete against rich institutions in recruiting the best students, postdocs, and beginning faculty. Clinician-researchers complain of severely cramped research space, despite their ability to attract greater external research funds and ICR dollars. Conversely, researchers in non-clinical units deplore their loss of state-derived hard-money salary, and feel UCSF focuses too much on clinical expansion to devote enough time or attention to research quality. In both groups, junior scientists focus on the excessive demands of grant-writing and San Francisco’s high cost of living.

## Choices for a sustainable future

UCSF and health centers like it must sustain themselves in an environment dramatically different from that of the 1980s. As clinical care became much more profitable, state support for academic missions greatly diminished, and many researchers found it difficult (at best) to obtain federal grants. UCSF’s leaders responded to the clinical sector’s profitability by embracing clinical expansion as a route to long-term survival in the health care marketplace, but no plan for responding to loss of research support has been defined. Although they profess admiration for ground-breaking research and its continuing support, these leaders—in sharp contrast to their predecessors 30 years ago—spend more time bragging about the national ranking of UCSF’s research revenues than thinking about scientific questions and quality or direction of research. They appear to assume that the institution’s research can easily take care of itself. “We do care about high-quality research!” exclaims one high official, adding: “Why do you think we build all these laboratories?”

Continuing reductions in external support make excellent research even more necessary. If the goal of a research university is to create new knowledge and understanding, leaders must not outsource their responsibility for research quality to external funders and donors. Can their institution afford to sustain higher quality? Which research goals and scope are most appropriate? UCSF sidesteps those hard questions by emphasizing quantitative assessments and promoting expansion. We believe UCSF cannot sustain its research mission—let alone its quality—without refocusing and selectively limiting that mission’s scope and revising its governance, funding, and administration. We present specific recommendations for UCSF, but some may prove useful for other publicly supported academic health centers.

### Research governance

We propose that all UCSF research be administratively and financially supported within an umbrella Division of Research (DOR). Such a division can help reverse existing fragmentation of the research mission in clinical departments and their divisions within clinical departments, and in basic science departments with little or no scientific rationale for existing as distinct academic silos. Silos impair scientific communication, resource sharing, and collaboration. Worse, fragmentation transforms the voices of individual silos into scattered chatter, easily ignored when the Campus makes a decision that affects all researchers. As shown below, DOR offers a mechanism for the necessary, difficult task of boosting research quality in both basic and clinical departments.

The DOR Director, a respected scientist who appreciates and understands clinical and basic investigation, will work with a Steering Committee of faculty researchers who represent research subdivisions within DOR—such as, perhaps, dry-lab, wet-lab, cell and molecular biology, genetics, and pathogenesis and treatment of disease. At the outset, DOR will augment, rather than supplant, existing departments and divisions. Using dollars derived from gifts, endowments, the Chancellor’s office, and the Schools, DOR will provide extra start-up dollars for recruiting outstanding junior faculty and for research supplements designed to mitigate the dangers of soft-money salaries. Search and promotion committees for faculty researchers will include members from DOR subdivisions and a relevant department or clinical division.

Because research comprises almost a quarter of all UCSF revenues, the DOR Director must belong to the highest leadership level, which now includes five individuals (12), none of whom is specifically responsible for the research mission. With enough philanthropic and other UCSF funds, and advice from the DOR Steering Committee, the Director can shape badly needed changes in direction and quality of UCSF research and ensure that decisions affecting research (e.g., for a new research building or project) reflect thoughtful input from UCSF’s researchers. Departments will retain their ability, in collaboration with DOR, to hire, promote, and support research faculty. Because some silo leaders will oppose DOR as a threat to their autonomy, the DOR approach will work only if Campus leaders strongly promote DOR’s ability to foster communication among departments and provide a needed voice in leadership decisions.

### Governance of basic science

DOR must tackle the critical task of re-shaping basic science research, which is imperiled by tough national competition, limited external funds, and inability to accumulate clinical dollars. UCSF’s present aspirations for fundamental biological investigation have not been articulated. Perhaps, as a basic science faculty member suggests, “to Campus leaders, basic science is a cute petting zoo that costs too much money and adds little value to the prosperous farm around it.” A key task for the DOR Director and the Campus will be hard-nosed financial assessment to determine whether or to what degree UCSF can support basic science. The assessment will probably show that UCSF resources can sustain long-term support for fundamental investigation only if the number of basic science faculty is reduced. To shrink that number gradually to a size that can sustain excellent research, supplemental DOR support will go only to the most outstanding present faculty, with the rest supported at a pre-DOR level; only a fraction of faculty retirements—perhaps two of every three—will open slots for recruiting new basic science faculty. New hires will be jointly chosen by DOR and the department, approved by DOR’s Steering Committee, and receive DOR support (startup costs, salary supplements, etc.). Basic science departments will be gradually replaced by a DOR subdivision, which will take responsibility for grant administration, salaries, promotion, and recruiting faculty. In this effort, the primacy of quality over quantity is critical: instead of employing vast numbers of researchers, the goal should be to retain and recruit a smaller number of very high-quality scientists with skills and knowledge broad enough for them to teach and learn from one another. They will collaborate both locally and within an electronically linked world-wide community.

### Philanthropy and a general endowment

Instead of letting quality fend for itself as UCSF research continues to grow without guidance, the DOR proposal provides a governance mechanism for making the necessary shift from quantity to quality. Such a shift will require leaders to demand high quality and make rather than avoid key fiscal decisions, especially with respect to philanthropy. Large gifts for research should not be accepted without deliberate institutional choices to use them to improve the overall research enterprise—a precept UCSF has been known to ignore (1).

Large private universities pay substantial fractions of operating costs from general endowments that grow every year. Astonishingly, despite steady loss of state support equivalent to billions of endowment dollars since the 1990s, UCSF has just begun to consider assembling a general endowment. At the outset such an endowment will be too small to make much difference, but failure to begin now will make a huge negative difference in 20 years—as, over the last 20, it already has. By rarely turning down gifts for building laboratories or endowments for any purpose, UCSF allows philanthropic donors to magnify the quantity-quality problem. Modest adjustments of that policy will require UCSF to favor gifts and a general endowment that broadly support its principal missions.

To accumulate a general endowment, UCSF can leverage its reputation for excellence in both clinical care and research, and persuade donors that general endowment dollars can promote excellence by populating its buildings with the very best clinicians and scientists, rather than trusting bricks and mortar to do the job. That excellence will require targeting more endowment dollars to outstanding faculty. The proposed general endowment would support investigators by paying salaries or research costs, from payouts administered by DOR, rather than departments or other units. Moreover, an increase—perhaps a doubling—of UCSF’s “gift assessment” tax on annual payouts from its directed endowments (1) could jump-start a general endowment, enhancing both fiscal viability and creative opportunities for faculty.

### Researcher salaries

Dependence on grants for large portions of faculty salaries saps loyalty to the institution and endangers creativity of researchers worried about their job security. Directing university dollars to individual researchers, to use either for salary or for research, will mitigate both effects. Instead of permanent professorships, UCSF should target dollars to faculty subgroups who need it, based on merit. For instance, UCSF’s Department of Medicine recently instituted a program for targeting $50,000 each year to faculty researchers, beginning with merit-based promotion to associate professor and continuing for a set period thereafter.

DOR and UCSF leaders should target support to subsets of researchers in a similar fashion. Consider these numbers: 5% annual payout ($12.5M) from a $250M endowment—close to the cost of its new neuroscience building—could have provided $70,000 per year to half the 358 associate professors who in 2014 received sponsored salary at UCSF (1). Awarded for any research purpose and based on merit judged by UCSF researchers, such extra dollars would enable chosen awardees to maximize research progress at a crucial time in their careers.

### Debt service

UCSF’s debt service obligation in 2025 will certainly exceed the $160M projected (see above). Along with costs of operating and populating new research facilities, this debt creates a continuing substantial drain on ICR, at present the main fungible source for supporting new projects and some portion of faculty salaries. For this reason, the DOR Director must carefully weigh costs and benefits of enhancing the quality of UCSF’s research enterprise against those of expanding it— that is, of investing in people *vs*. bricks. Rather than passively acquiesce to blind expansionism and donors’ passions for buildings, the Campus should choose among possible directions for its research. A UCSF capital campaign should commit future philanthropic dollars to a general endowment, establish a DOR, reduce the rate at which debt and debt service rise, and support the very best scientists.

## Can UCSF deliberately determine the future of its research?

For UCSF and similar academic centers, this is a crucial question. Deliberate change at UCSF will be powerfully opposed by economic, social, and political forces that buffet the institution from outside and by the inertial forces of its internal governance, culture, leaders, faculty, and staff. Still, UCSF has already accomplished two successful and deliberate major changes: in the 1970s and 1980s, it transformed itself from an ordinary state-supported health center into a research powerhouse (13); in the 1990s, it expanded and improved the quality of its clinical enterprise (1). Both changes were aided by external forces—fast-growing NIH budgets and the DNA revolution in one case, exuberant growth of the medical care industry in the other. In contrast, our proposed changes run directly counter to formidable forces within and outside UCSF, including clinical expansion and the corollary idea that more research is always better. We fear that leaders and faculty will choose to ignore not only corrective proposals, but also the problems they try to correct, relying on clinical expansion to save the day and leaving the entire research mission—like the basic science petting zoo—to wage battle alone. In doing so, they will cast aside both financial prudence and the university’s academic duty to create new knowledge.

Prudence first: to preserve research as its support diminishes, it is foolish to build more facilities, borrow more money, and hire more scientists to compete for ever-scarcer and more arbitrary NIH awards. Rather than expand capacity, it is prudent to use gifts and endowments to ensure security and quality of a financially jeopardized mission. To do so, UCSF must value high-quality research as the lifeblood of a research university, recognize that readily accessible government support will not increase soon, and resist the national delusion that academic research can pay all its costs. Prudence also dictates attempts to replace the generous research support UCSF once enjoyed, from private (non-government) sources and philanthropy. Because private industrial support goes preferentially to clinically oriented research, the university will have to decide whether philanthropy can support its fundamental research at a level sufficient to enhance its quality. If not, UCSF should gradually shrink basic research and focus primarily on more readily supported clinically-driven research.

As for the academic duty to create new knowledge, consider Clark Kerr’s prediction, in 1963, that knowledge produced by research universities would enhance economic prosperity and security of a growing nation after World War II (14). Five decades later, the benefits of biomedical research prove Kerr’s case. New knowledge generated by academic science produces rapid changes in our world, explains them, and helps us respond to them. Now, even more than in the 1960s, the world badly needs continuing creative scientific discovery from academic research. Despite economic and social forces that reduce research revenues, UCSF and other publicly supported universities must continue to produce the new knowledge necessary to meet future challenges.

